# Back to the future: Implications of genetic complexity for hybrid breeding strategies

**DOI:** 10.1101/2020.10.21.349332

**Authors:** Frank Technow, Dean Podlich, Mark Cooper

## Abstract

Commercial hybrid breeding operations can be described as decentralized networks of smaller, more or less isolated breeding programs. There is further a tendency for the disproportionate use of successful inbred lines for generating the next generation of recombinants, which has led to a series of significant bottlenecks, particularly in the history of the North American and European maize germplasm. Both the decentralization and the disproportionate inbred use reduce effective population size and constrain the accessible genetic space. Under these conditions, long term response to selection is not expected to be optimal under the classical infinitesimal model of quantitative genetics. In this study we therefore aim to propose an alternative rational for the success of large breeding operations in the context of genetic complexity arising from the structure and properties of interactive genetic networks. For this we use simulations based on the *NK* model of genetic architecture. We indeed found that constraining genetic space and reducing effective population size, through program decentralization and disproportionate inbred use, is required to expose additive genetic variation and thus facilitate heritable genetic gains. These results introduce new insights into why the historically grown structure of hybrid breeding programs was successful in improving the yield potential of hybrid crops over the last century. We also hope that a renewed appreciation for “why things worked” in the past can guide the adoption of novel technologies and the design of future breeding strategies for navigating biological complexity.

## Introduction

Pioneered by Shull (1908), hybrid breeding is credited as one of the most significant factors for the tremendous productivity increases of major field (Duvick, 1999) and horticultural (Silva Dias, 2010) crops that enabled food production to keep pace with population growth. Hybrid breeding programs originally were centred around maximum exploitation of heterosis, a phenomenon that remains largely unexplained even after a century of research (East, 1936; Lippman and Zamir, 2007). This later evolved into the modern concept of hybrid breeding, characterized by its distinctive structuring of germplasm into heterotic groups and patterns (Melchinger and Gumber, 1998). Beyond heterotic groups, the structure of commercial hybrid breeding, particularly in major crops like maize, is characterized by the largely isolated and unique sub-heterotic patterns of the major companies (White et al., 2020) as well as a high degree of decentralization into smaller, more or less disconnected sub-programs within those (Cooper et al., 2014). Plant breeders further have a tendency for relying on only a small set of elite inbred lines for producing the next generation of recombinants (Rasmusson and Phillips, 1997), leading to a series of significant bottleneck events in the history of, for example, the North American maize germplasm (White et al., 2020). These characteristics drastically reduced the effective population size within breeding programs and are not predicted to be promising strategies under the additive, infinitesimal model of quantitative genetics (Gaynor et al., 2017). Nevertheless, consistent long-term genetic gain has been demonstrated (Duvick et al., 2004).

To better describe and quantify the observed genetic variation among hybrids, the concept of general and specific combining ability was developed early on (Sprague and Tatum, 1942). The former, commonly abbreviated as GCA, is a property of the additive effects of contributing genes and describes the average performance of all hybrids derived from an inbred. The latter, commonly abbreviated as SCA, is a non-additive residual term that describes the deviation of the performance of a particular hybrid from the expectation based on the parental GCA values.

Running efficient hybrid breeding programs requires a preponderance of additive genetic variation to maximize response to selection in the next generation of inbred lines (Falconer and Mackay, 1996) as well as the predictability of hybrid performance from the GCA of inbred lines (Reif et al., 2007). A preponderance of GCA variation also allows identification of inbreds that can serve as parents of several high performing hybrids. This greatly simplifies production of commercial seed, which is a major challenge for many crops (Technow, 2019). Therefore, hybrid breeding programs have traditionally relied on maximizing and exploiting GCA variation (Falconer and Mackay, 1996; Melchinger, 1999).

The historically grown paradigms around hybrid breeding designs and strategies are now being challenged by innovative concepts (e.g., Gaynor et al., 2017; Wallace et al., 2018; Hickey et al., 2019; Voss-Fels et al., 2019; Seye et al., 2020) devised in the wake of technological advances such as whole genome prediction (Meuwissen et al., 2001), high-throughput phenotyping (Araus and Cairns, 2014) and genotyping (Poland and Rife, 2012), as well as gene editing (Jaganathan et al., 2018). While some of these concepts are highly speculative and might not live up to expectations (Bernardo, 2016), it is clear that the next decades will change plant breeding. However, before implementing drastic changes to breeding programs we require a theoretical and simulation framework to explore and understand the structures and strategies that have contributed to the success of long term genetic gain and germplasm improvement. From this historical basis we can evaluate novel proposals and draw lessons for design of future breeding strategies.

Empirical reports show a preponderance of additive variation in many wild, domesticated and laboratory species (Falconer and Mackay, 1996; Lynch and Walsh, 1998; Hill et al., 2008). This agrees well with published studies showing a preponderance of additive GCA over non-additive SCA variation in hybrid breeding programs (Technow et al., 2014; Larièpe et al., 2017). At the same time, however, advances in plant physiology and molecular and systems biology have stimulated a renewed appreciation of the intricate interactions at the molecular, metabolical and physiological level that underlie complex traits (Hammer et al., 2006; Carlborg and Haley, 2004; Phillips, 2008; Wilkins et al., 2016; Saha et al., 2011; Jiang et al., 2017). Of particular relevance for hybrid breeding are recent studies indicating that heterosis is an emergent property of complex metabolic networks (Fiévet et al., 2010, 2018; Vacher and Small, 2019).

The paradox between the complexity of the underlying biology and the simplicity of the expressed variation can of course be resolved by distinguishing between biological and statistical effects and realizing that the former cannot be inferred from the latter (Wade, 2002; Mackay, 2014; Huang and Mackay, 2016). Statistical effects of genes, as well as their aggregates such as GCA and SCA and their variances depend on the genetic background of the population in which they are evaluated, particularly on allele frequencies and linkage disequilibrium (LD) patterns (Falconer and Mackay, 1996). For example, it was shown that, regardless of the underlying genetic architecture, genetic variances in random mating populations are expected to be predominantly additive when genes are at extreme frequencies and linkage disequilibrium is high (Hill et al., 2008).

Thus, ratios of additive to non-additive variation are not intrinsic properties of biological systems but at least partly a function of allele frequencies and LD patterns and thus dependent on breeding strategies. Because of the importance of additive variation for efficient operation of breeding programs, a framework to evaluate and study breeding strategies should allow for the possibility of additivity arising from high degrees of biological complexity at the genetic level.

In this study we will use simulations based on the *NK* model of genetic complexity (Kauffman, 1993) to explore two *themes* representing key, historically grown, characteristics of hybrid breeding: firstly its decentralization into smaller, more or less independent sub-programs (“*decentrality* theme”) and secondly the disproportional use of superior inbred lines for producing the next generation of recombinants (“*inbred usage* theme”). Our goal thereby is not to make specific recommendations for optimal structuring of programs, but rather to gain an appreciation for the properties of these structures in the context of different degrees of genetic complexity.

## Material and Methods

### Model of genetic complexity

The *NK* model, introduced by Kauffman (1993) will form the basis of the simulations. The *NK* model allows generation of a tunable series of models of trait genetic architecture with increasing dimensionality and complexity by varying the number of genes *N* (dimensionality) and the degree of interaction among them (*K*, complexity).

The *genetic landscape* metaphor was introduced and developed by Wright (1932) to aid conceptualizing genetic complexity in high dimensions. As a metaphor it should not be taken literally but can help to gain an intuition for the complexity and ruggedness associated with increasing values of *K* (Kauffman, 1993) as well as making the rather abstract concepts discussed henceforth more tangible. At *K* = 1 (special case of additive gene action), the genetic landscape can be imagined as that of Mount Fuji, i.e., a single, clearly distinguished peak with a steady and monotonous incline to the top (Figure 1). At intermediate *K* levels, the landscape is characterized by multiple peaks clustered together in a certain region of genetic space. This might be visualized as akin to the European Alps, i.e., a mountainous region within an otherwise flat landscape. Finally, at high value of *K*, the landscape resembles a sea of dunes, i.e., a range of peaks of similar height and shape distributed more or less evenly in space.

**Figure 1.**
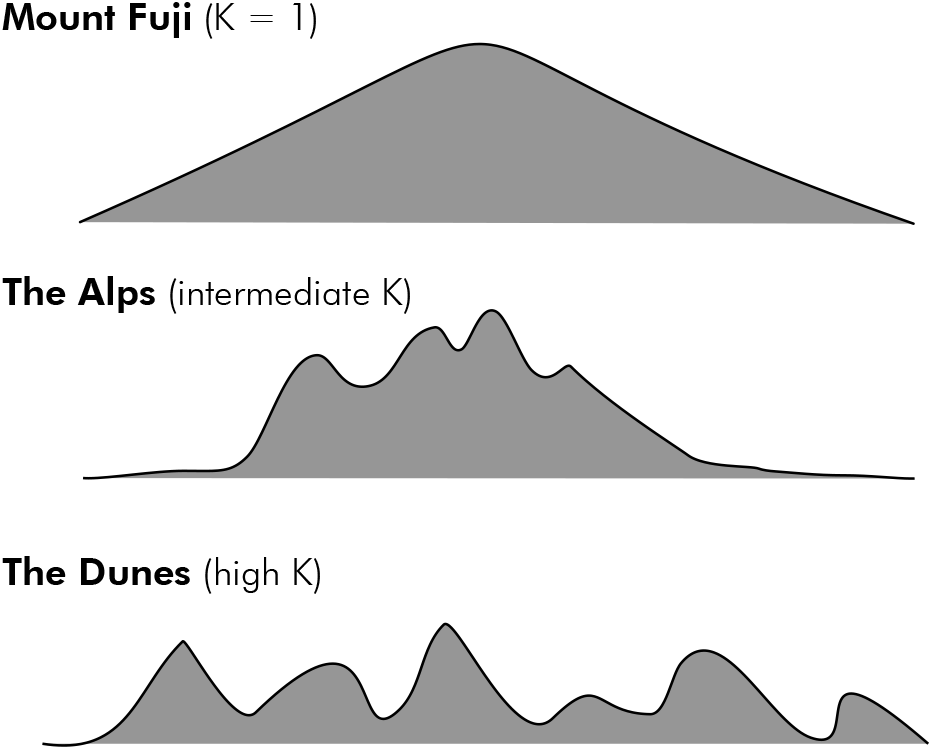
Schematic visualization of genetic landscapes corresponding to different values of complexity parameter *K*.

We implemented the *NK* model according to the generalized approach described by Altenberg (1994), but adapted the model to accommodate diploid genomes. Here, the complex trait is described as a normalized sum of a set of “fitness components”. The value of each fitness component is computed as a function of *K* interacting genes drawn at random from all *N* genes. Following Altenberg (1994), the specific fitness values were calculated with *random functions* derived from the *ran4* pseudo-random number generator (Press et al., 1992) and are distributed uniformly between 0 and 1. For this study, the number of fitness components and the number of genes were both set to 500. Genes were biallelic and the simulated organism diploid. The complexity parameter *K* was varied from 1 to 15 in steps of 1 (i.e. creating genetic landscapes ranging in complexity from Mount Fuji to The Dunes; Figure 1). With the exception of *K* = 1, this parameter was used as the rate parameter in a Poisson distribution from which the number of interacting genes was drawn independently for each fitness component. The sampled values were then truncated to fall within a range of 1 and 15. The identities of the interacting genes were drawn at random from the total set. Thus genes typically influenced multiple fitness components (i.e., act pleiotropically). For *K* = 1, each of the 500 genes was assigned to exactly one fitness component and the values of heterozygous allele configurations constrained to be midway between the homozygous configurations. Thus, *K* = 1 represents the special case of additive gene action. Using order statistics, the expected value of the maximum of two samples from a Uniform distribution between 0 and 1 is 2/3. Thus, the expected maximum attainable fitness for the *K* = 1 special case is 2/3.

The complexity of the generated *NK* models was quantified following the “one-mutant neighbour” hill-climbing algorithm described by Kauffman (1993), but adapted to diploid organisms. A randomly generated genotype was used as the starting value. From there, all possible genotypes were generated that differ from the initial genotype by one allele at one of the 500 loci. Thus, a homozygous locus was changed to the heterozygous state while a heterozygous locus was changed to both alternate homozygotes. Then, the fitness values of all one-allele neighbours were evaluated according to the defined *NK* model and an improved genotype chosen at random from all fitter one-allele neighbours. This process was repeated until no fitter one-mutant neighbour could be found, meaning that the search reached a local or global optimum. For each level of *K*, 100 *NK* models were generated independently and a minimum of 65 searches, each starting at a random initial genotype, were conducted for each. The statistics recorded were the average number of steps until a local optimum was reached, the average Hamming genotypic distance, i.e., the normalized number of differing genome positions (Pinheiro et al., 2005), among optima and the correlation between the fitness values of the optima and the Hamming distance to the highest optimum identified (Kauffman, 1993).

The average Hamming distance between the local peaks increased from just below 0.5 at *K* = 2 to 2/3 at around *K* of 6 or 7 and remained constant at this value from there on (Figure 2A). Note that with three different genotypes at each locus, 2/3 is the expected value of the Hamming distance between randomly generated genotypes. Similarly, the correlation between the fitness values of the local peaks and their Hamming distance to the highest identified peak increased from −0.25 at *K* = 2 to zero at *K* = 9 (Figure 2B). Here, a negative correlation means that local peaks with higher fitness tend to be found near each other and clustered around the highest peak. Further, a zero correlation indicates that there is no clustering of the peaks and proximity to the highest peak. Therefore, local peaks are randomly distributed throughout the genetic landscape. Thus, somewhere between *K* = 6 and *K* = 9, the landscape shifts from one in which local peaks tend to cluster together, to one where local peaks of arbitrary height can exist anywhere in genetic space. The average number of steps until a local peak was reached decreased with *K* from 500 at *K* = 1 to just 167 at *K* = 15 (Figure 2C). Note that 500 is the expectation at *K* = 1, the special case of additive gene action, when starting from randomly generated genotypes, because 1/3 of the 500 loci are already at their highest possible value, 1/3 are one step removed (the heterozygous genotypes) and 1/3 are two steps removed (the lower homozygotes). Thus, the complexity and ruggedness of the genetic landscapes increase further after they become uncorrelated around *K* of 6 to 9.

**Figure 2.**
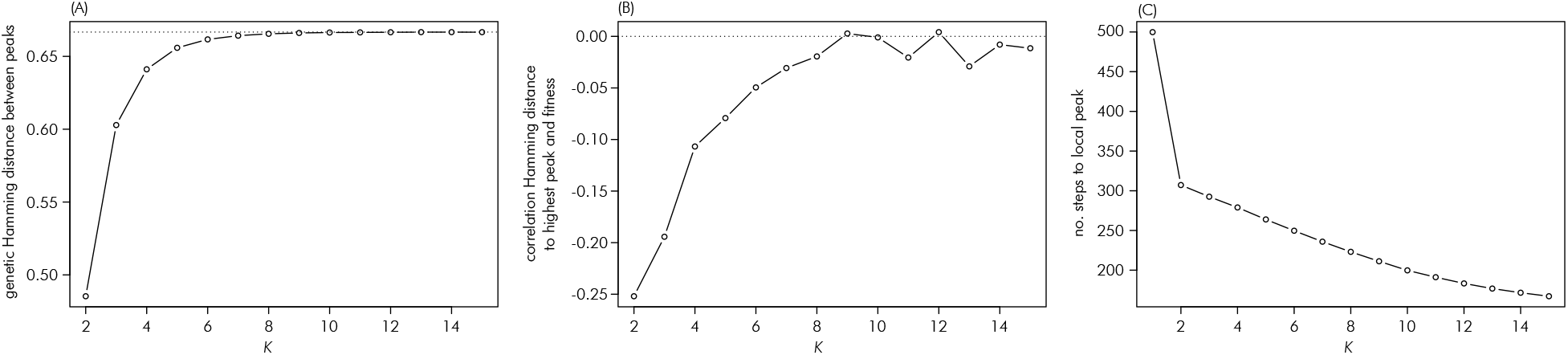
Relationship between the *NK* model complexity parameter *K* and (A) the average genetic Hamming distance between local peaks, (B) the average correlation between the fitness values of local peaks and their genetic Hamming distance to the highest peak and (C) the average number of steps to a local peak. Panels (A) and (B) omit results for *K* = 1, for which only a single peak exists. The dotted lines in these panels indicate values of 2/3 and 0.0, respectively.

The simulated genome comprised 10 diploid chromosomes of 1 Morgan length each. Each of the chromosomes received a random subset of 50 of the 500 genes, which were distributed evenly across the chromosome. Recombination was simulated according to the Haldane mapping function with the R package “hypred” (Technow, 2013), in the version available from the supplement of Technow and Gerke (2017).

### Simulation of hybrid breeding process

The simulation process is visualized in Figure 3. The starting point of the simulation was a base population of inbred lines of size 1,000. This population was simulated stochastically as described by Montana (2005) to result in an expected LD between two loci *t* Morgan apart equal to *r*^2^ = 0.5 · 2^−*t*/0.1^ and with minor allele frequencies distributed uniformly between 0.35 and 0.50. The lines from the base population were then separated at random into two heterotic groups (arbitrarily labelled ‘1’ and ‘2’) and further into sub-populations within those. The size of those sub-populations depended on the scenario. One sub-population from one heterotic group was then paired with one sub-population from the other group to form sub-heterotic patterns. These population pairs will henceforth be referred to as “breeding programs”. Hybrids were produced strictly across heterotic groups, by crossing lines from one sub-population of a program with lines from the other. Breeding crosses, i.e., crosses to generate a new generation of recombinant lines were done within and among sub-populations, depending on the scenario but strictly within heterotic groups. The simulation of the breeding process described above is an approximation of the structure and evolution of long term hybrid breeding akin to what we have observed in practice; i.e. starting from an initial germplasm base, separation into distinct heterotic groups and future separation into sub-populations.

**Figure 3.**
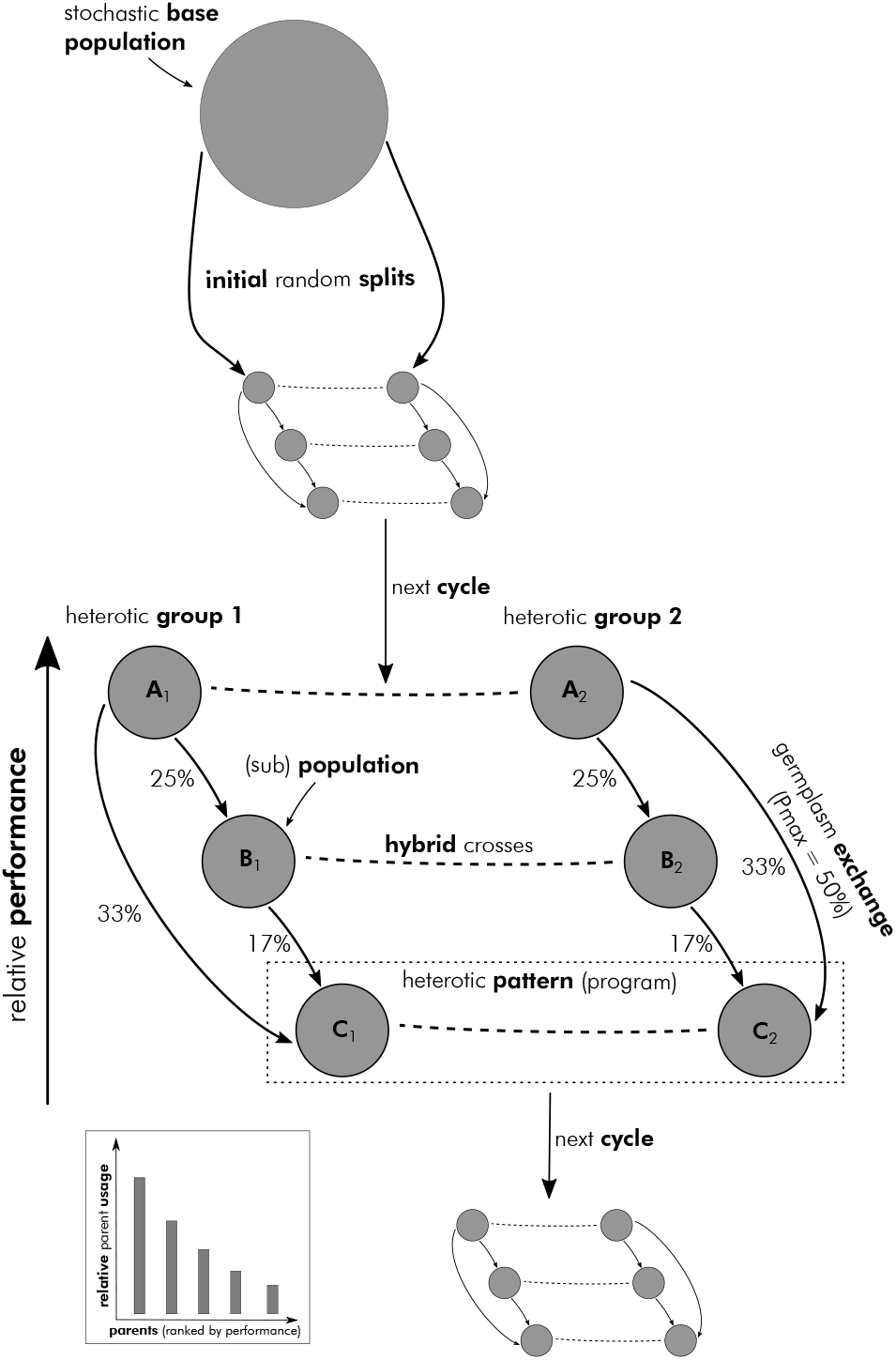
Schematic visualization of simulated hybrid breeding process.

The GCA of the lines was evaluated with an incomplete mating design (Melchinger et al., 1987; Seye et al., 2020) by performing 10 crosses per line with random partners from the opposite sub-population of the same program. The performance of the resulting hybrids, as determined according to the defined *NK* model, was then averaged. Finally, a normally distributed noise variable with zero mean and variance equal to one third of the variance of the GCA values of that sub-population was added to those averages to represent experimental and environmental noise. The so obtained values were used as observed GCA values. Those GCA estimates were then used to predict the performance of all possible inter-group hybrids of that program. The top hybrids, how many exactly depended on the scenario, were then selected and their true performance determined according to the *NK* model. The average of this select group of hybrids, which represents a set of advanced experimental hybrids, was used to quantify the overall performance of the program in the current cycle. The maximum true performance of the selected hybrids from all programs was defined as the peak performance of the whole breeding operation in the current cycle and used as a metric of genetic gain. This metric reflects that commercial breeding programs release only a handful of hybrid products each cycle.

Breeding crosses among inbred lines for initiating the next recombination cycle were chosen by assigning each inbred line a usage probability, which was a product between an individual and population level relative contribution value. To determine the former, the lines within each sub-population were ranked according to their observed GCA values. Only the top lines, how many depended on the scenario, were selected as potential parents, the remainder given an individual contribution value of zero. The relative contributions of the selected lines from a given sub-population were drawn from a Dirichlet distribution. The concentration parameters of this distribution were used to modulate the relationship between selection rank and relative contribution. Further details about this will be given later when describing the setting for the “inbred usage theme”.

The population level contribution values describe the overall contribution of lines from one sub-population to the breeding crosses of another. They are thus defined anew for each target population and hence the contribution value of population ‘A’ to the crosses for population ‘B’ might be different than that to the crosses of population ‘C’. The process will be explained using the example visualized in Figure 3. Here there are three programs (labelled ‘A’, ‘B’ and ‘C’, with subscript 1 or 2 indicating the heterotic group).

The programs are ranked from highest to lowest performing according to the average performance of the selected set of experimental hybrids, as described above. Germplasm, in the form of lines used as crossing partners, is exchanged only from higher performing to lower performing programs. Specifically, the amount of crosses with lines from other programs increased from zero for the best performing program (A) to a proportion of *P*_*max*_ for the lowest performing program (C), with intermediate programs staggered equidistantly between. In the example, *P*_*max*_ = 50%. Thus, program A will perform no crosses with lines from other programs, program B will use lines from other programs in 25% of its new crosses and program C in 50% of its crosses. How much of that overall proportion was derived from each of the other programs was proportional to the relative performance differences. In the example, the difference between program C and A is twice as large as that between C and B, thus, lines from program A were used in twice as many crosses than lines from program B (33% from A and 17% from B for at total of 50%). This process thus reflects that highly successful programs tend to exploit their own genetics while less successful programs have more of an incentive to explore superior genetics from other programs.

The relative individual contributions were then multiplied with the relative population contributions to arrive at a final relative contribution value for each line to the crosses of a given population. The actual breeding crosses were then determined by sampling the lines with probabilities proportional to their contribution values. This was done with replacement, meaning that the same cross could have been made multiple times, but excluding crosses that would result in selfings. One recombinant line was derived from each crossing through seven generations of single seed descent selfing, followed by a final doubled haploid step (Dwivedi et al., 2015) to result in fully homozygous inbred lines. This new generation of recombinants fully replaced the previous generations, i.e., a line was considered as a crossing partner in only one generation. The so obtained new recombinants then form the next breeding cycle. The simulations were conducted for 30 cycles in total and repeated independently at least 500 times for each scenario studied. All computations were conducted in the R environment for statistical computing (R Core Team, 2018).

### Recorded metrics

In addition to the already described true performance of the best identified hybrid, which was used as a measure of *peak performance* in a given cycle, several other measures were recorded to describe and understand the dynamics of the system.

The *proportion of GCA to total genetic variance* (%GCA) describes the amount of exploitable additive genetic variation currently available. It was estimated using the hybrids generated for evaluating the GCA of the inbred lines. For this, the following mixed model was fitted: *h*_*ij*_ = *μ* + *g*_*i*_ + *g*_*j*_ + *e*_*ij*_, where *h*_*ij*_ was the true performance of the hybrids, *μ* the overal mean, *g*_*i*_ and *g*_*j*_ were the GCA effects of the parents from the two heterotic groups and *e*_*ij*_ a residual term. Because the true genetic performances of the hybrids were used, *e*_*ij*_ corresponds to the SCA component. The model was fitted using the R package “lme4” (Bates et al., 2015) and %GCA then calculated as (*V*_*gi*_ + *V*_*gj*_) / (*V*_*gi*_ + *V*_*gj*_ + *V*_*eij*_), where *V*_*gi*_ etc. were the estimated variance components. In scenarios with multiple programs, %GCA was estimated separately for each and then averaged to arrive at a single estimate for each cycle.

The *modified Rogers’ distances* (Reif et al., 2005) between the heterotic groups within each program were used as measures of heterotic group divergence. The distances were calculated for all programs and averaged to arrive at representative value for that cycle.

To describe the distribution of allele frequencies within each sub-population and hence the amount of available allelic diversity we calculated the proportion of loci with a minor allele frequency of less than 5%. This probability measures the thickness of the extreme tail of the allele frequency distribution and thus reflects the degree with which it follows a ‘*U-shape* ’ (Hill et al., 2008). This metric was evaluated for all sub-populations in each cycle and then averaged.

As a more high-level diversity metric we considered the *effective population size* (*N*_*e*_) of each sub-population. *N*_*e*_ was calculated according to the method described by Corbin et al. (2012) for estimating constant effective population size. The so obtained values were averaged across sub-populations.

### Hybrid breeding ‘themes’

All previously described parameters, such as parameters related to the *NK* model and genetic architecture, parameters related to testcross evaluation, etc, were kept constant across the themes investigated.

In the *decentrality* theme we explored consequences of separating hybrid breeding programs into smaller, more or less isolated, units. We defined three distinct strategies for ‘searching’ (Podlich and Cooper, 1999) genetic space (Figure 4): a single large program (*centralized search*) to multiple smaller, fully isolated programs (*isolated search*). Between these two extremes we considered a strategy with multiple smaller programs that exchange germplasm in the form of breeding crosses (*distributed search*).

**Figure 4.**
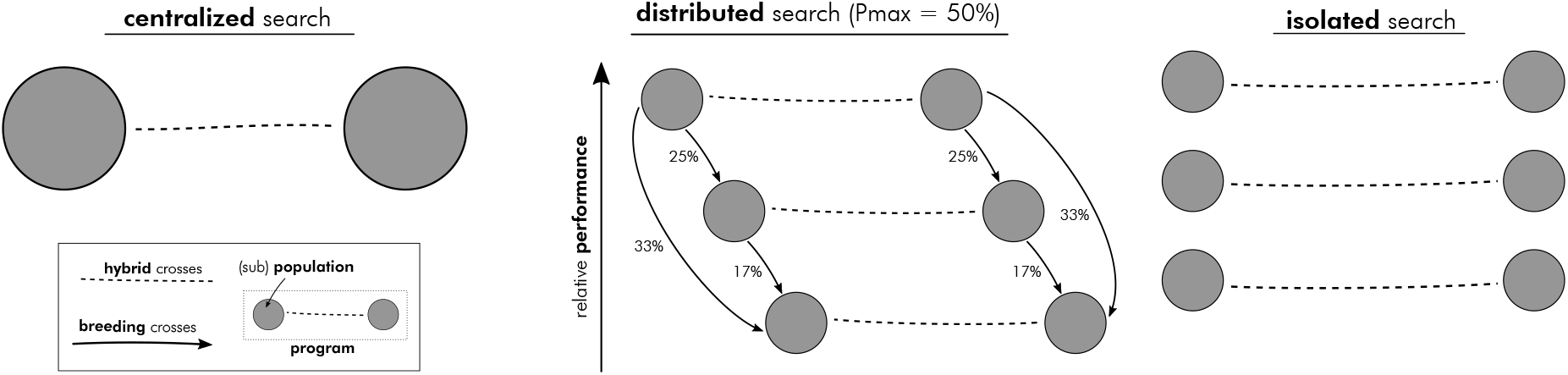
Schematic visualization of the three general search strategies explored in the *decentrality* theme.

The centralized search was characterized by a single program consisting of one sub-population per heterotic group. The size of each was 500, for a total of 1,000 lines generated in each cycle. The number of lines selected to contribute to the next generation was 125 per sub-population. The relative individual contributions of these lines decreased proportionally with their performance ranks. The number of selected experimental hybrids was 125. The isolated search strategy comprised five programs, each with one sub-population per heterotic group. The sub-population size was 100 of which 25 were selected. Also here, the relative individual contributions of the lines were proportional to their performance ranks. The number of experimental hybrids selected per program was 25.

In the distributed search we considered three levels of Pmax: 25%, 50% and 75%. The number of programs as well as lines and hybrids created and selected for each followed those of the isolated search. Note, thus, that the total number of lines and hybrids were the same across all scenarios as was the selection intensity.

In the *inbred usage* theme we explored the consequences of different degrees of imbalance in the relative contributions of the selected inbred lines to the next generation. The different scenarios explored correspond to the distributed search strategy with Pmax = 50%. Only the relative usage of inbred lines was varied. As described above, the observed relative contributions were drawn from a Dirichlet distribution with concentration parameter chosen in a way to result in a certain average relationship between relative contribution and performance rank. Three scenarios were considered (Figure 5). In the *balanced* usage scenario, all selected inbreds contributed equally on average, in the *proportional* scenario, the relative contribution declined proportional with the performance rank of the lines. In the *disproportional* scenario, contributions halved with every 5 ranks, meaning that the highest performing line will contribute twice as much to the next generation as the 5th ranked line. The increasing imbalance in contributions can be quantified as 1/***b′b*** (with ***b*** being the vector of relative contributions), which is an estimate of the effective number of contributing lines (Boichard et al., 1997). For the balanced scenario, this was 25 and thus equal to the actual number of selected lines within each sub-population. It decreased to 19.1 and 13.6 for the proportional and disproportional scenarios, respectively.

**Figure 5.**
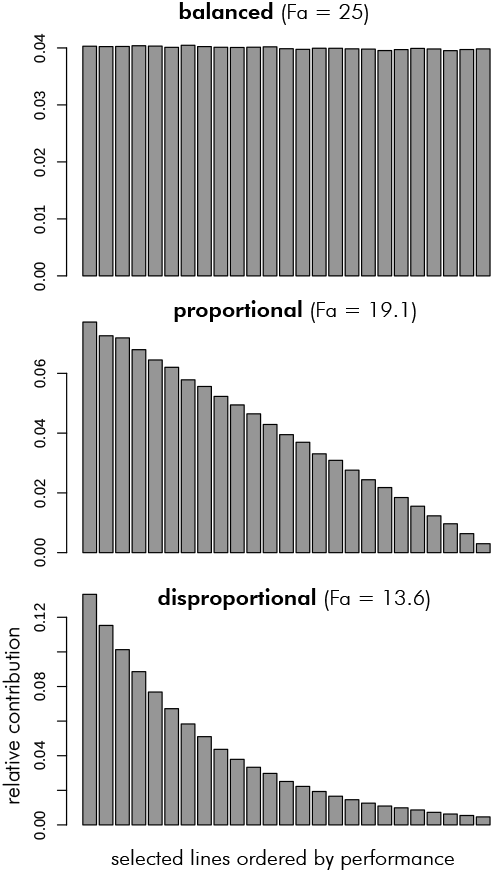
Distributions of relative individual contributions of selected inbred lines considered in the *inbred usage* theme.

## Results

### Decentrality theme

Which strategy achieved the highest peak performance depended on the value of the complexity parameter *K*, with the centralized strategy being superior at low *K* < 5, the distributed strategy at intermediate *K* and the isolated strategy at high values of *K* above eight (Figure 6A). The differences between the strategies tended to increase with increasing *K*. The centralized and distributed search strategies came very close to reaching the theoretical maximum peak performance at the special case of additivity (*K* = 1), but the isolated strategy remained considerably below that. Within the distributed strategy, the highest Pmax value of 75% was superior at *K* values below eight and the lowest Pmax of 25% at high *K* (Figure 6B). The case of Pmax = 50% had peak performance in between the two extremes, but more similar to Pmax = 75%. All Pmax scenarios achieved virtually identical peak performance at *K* = 1.

**Figure 6.**
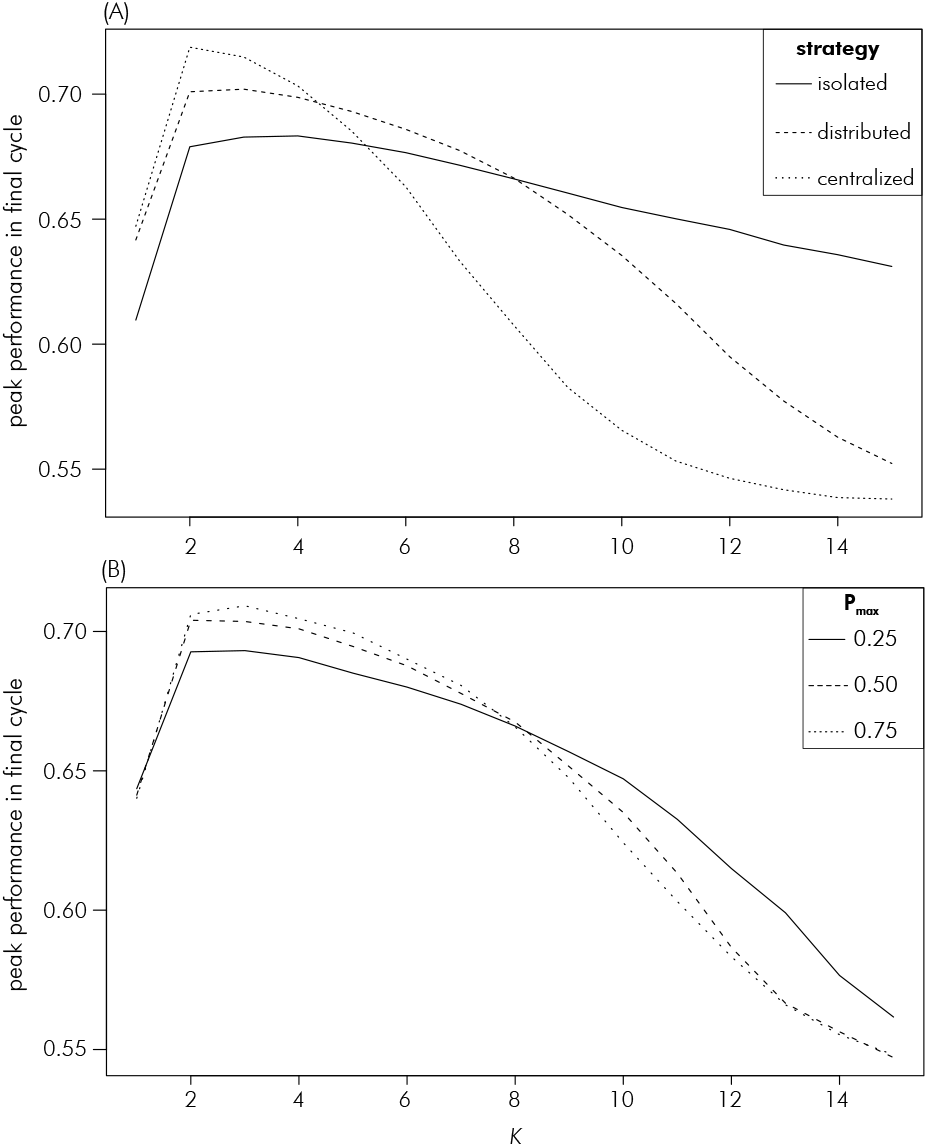
Relationship between the *NK* model complexity parameter *K* (average number of interacting genes) and peak genetic performance in the last cycle for the strategies explored in the decentrality theme: (A) comparing the isolated, distributed and centralized strategies and (B) the different values of Pmax within the distributed strategy. The curve of the distributed strategy in (A) is an average across the three Pmax scenarios within it.

For brevity, trajectories across cycles are shown only for *K* values of 1, 6 and 15, representing the additive, multi-peaked but clustered and fully uncorrelated landscapes, respectively (Figure 1). Results for all values of *K* are available as supplemental information (File S1). At *K* = 1, the centralized search strategy had the highest peak performance in all cycles, closely followed by the three versions of the distributed search (Figures 8A, B, C). The peak performance of the isolated search strategy was considerably lower than that of the other strategies as it increased at a lower rate and seemed to reach a plateau at around cycle 20. At *K* = 6, the isolated search strategy achieved the highest peak performances in the earlier cycles but was overtaken by the distributed search strategies later. Those had very similar peak performances until the last few cycles when the version with Pmax of 25% fell behind. The centralized search had the lowest peak performances throughout, with the differences to the other strategies being particularly large between the intermediate cycles 15–20. Finally, at *K* = 15, only the isolated search had a sizable increase in peak performance cycle over cycle. The distributed search strategies showed an increase only in the last few cycles and the centralized strategy did not increase peak performance at all.

As expected %GCA was equal to one for all scenarios at *K* = 1 (Figures 8D, E, F). At *K* = 6, %GCA started at just below 10% and increased from there with each cycle. The rate of increase was greatest for the isolated strategy which reached almost 100% in the final cycles. The centralized search strategy had the slowest increase and was still below 50% in the final cycle. The distributed search strategies were intermediate between these two extremes. The increase was steepest for Pmax = 25% case, which translated to it having a markedly higher %GCA than the Pmax = 50% case and Pmax = 75% case during intermediate cycles 15–20. However, all three converged to a similar value of around 80% in the final cycle. At *K* = 15, %GCA started at zero and only the isolated strategy saw a marked increase in early cycles. The distributed search strategies saw an increase in %GCA noticeably above zero only in the final cycles and the centralized strategy remained at zero throughout.

The percent of loci with MAF < 0.05 increased over cycles for all strategies and complexity levels (Figures 8G, H, I). In all cases, the increase over cycles was strongest for the isolated search strategy, where it reached close to 100% in the final cycles and weakest in the centralized search strategy. The curves for the three Pmax levels of the distributed search strategy were similar to each other and intermediate compared to the two other strategies. The differences between the strategies increased with *K* because the increase in the proportion of loci at extreme frequencies slowed for the distributed and centralized strategies with increasing *K*. At the highest levels of complexity, only between 30% and 40% of loci showed a MAF < 0.05 in the different distributed strategies and less than 20% in the centralized strategy.

The modified Rogers Distance between heterotic groups increased over cycles for all strategies (Figures 8J, K, L). For all levels of complexity, this distance was highest for the isolated strategy and lowest for the centralized strategy, with the three versions of the distributed search having similar values that were intermediate to the two extremes (Figure 7).

**Figure 7.**
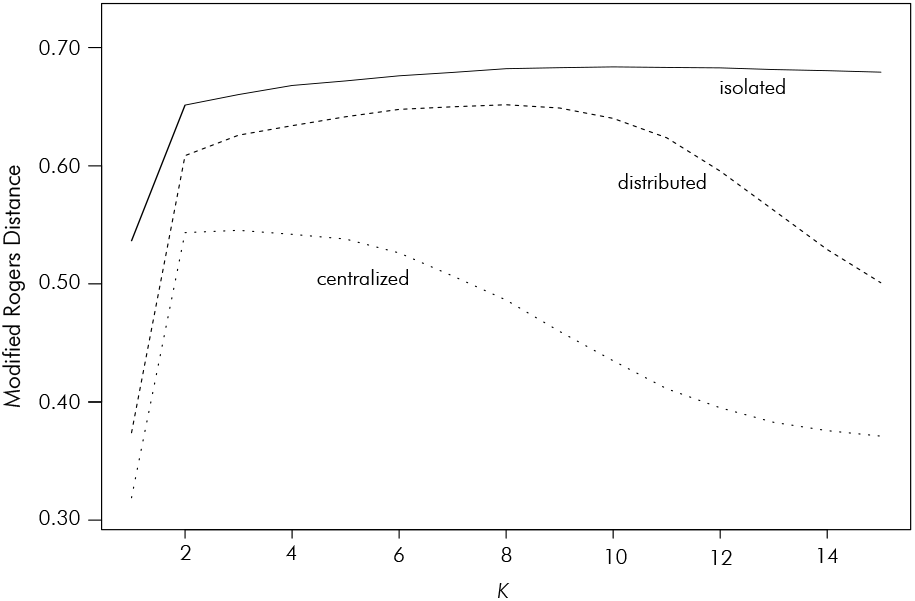
Modified Rogers Distance between heterotic groups as a function *K*, averaged across programs, in the connectivity theme. The curve of the distributed strategy represents the average across the three Pmax levels.

**Figure 8.**
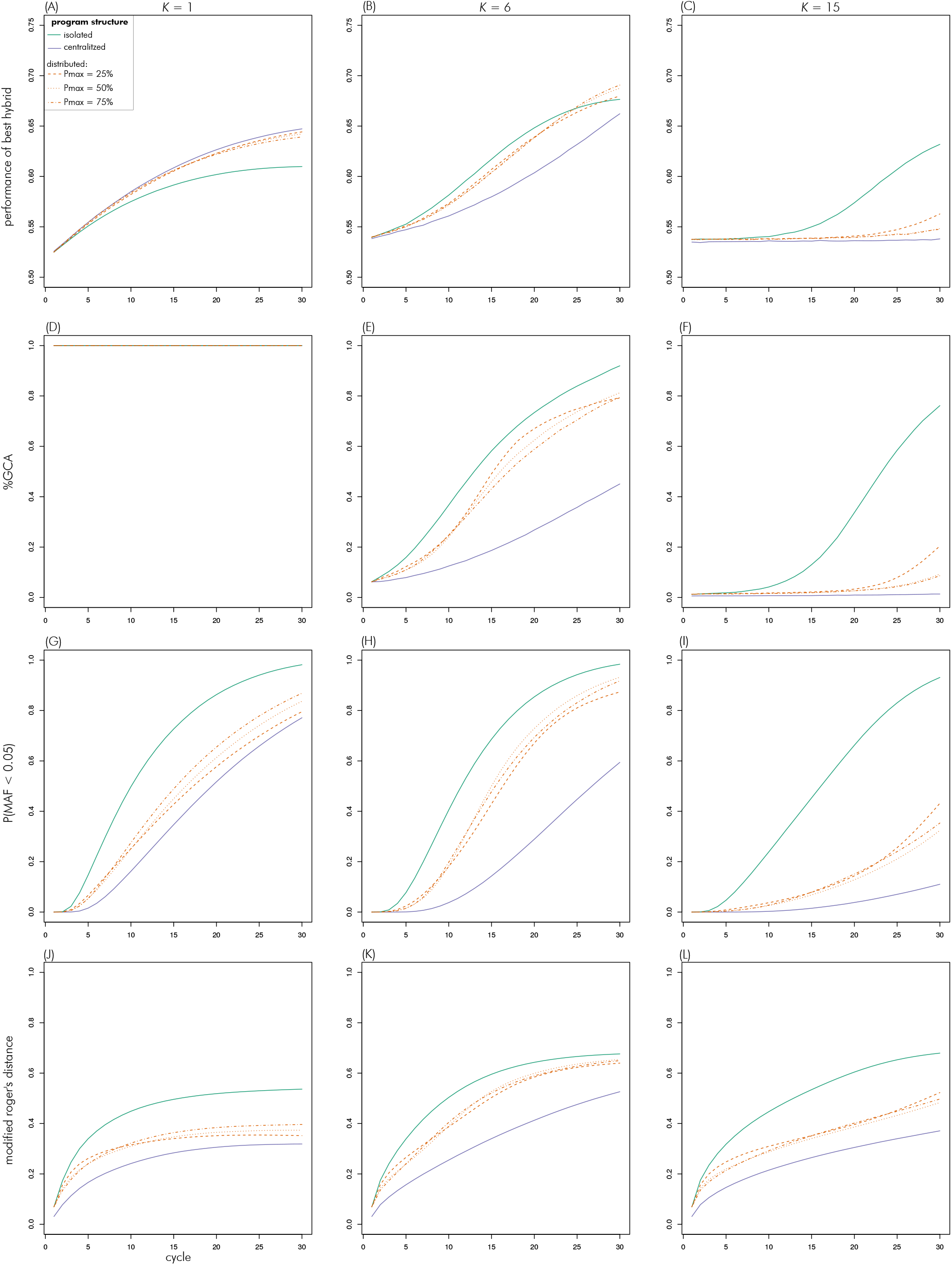
Evolution of metrics over cycles in the decentrality theme for scenarios with *K* of 1 (left column), 6 (middle) and 15 (right).

The *N*_*e*_ differences among the strategies remained largely constant across cycles and levels of *K*. For the sake of brevity results will only be reported for cycle 15 and *K* = 7. The estimated *N*_*e*_ for each-sub-population for the isolated search strategy was 20.0, for the three versions of the distributed search it was 23.7 (Pmax = 25%), 31.3 (Pmax = 50%) and 35.4 (Pmax = 75%), respectively, and for the centralized search strategy 98.3.

### Inbred usage theme

For brevity sake, only results for *K* of 1, 6 and 15 are shown (results for all values of *K* are provided in supplemental file S2). Again, which inbred usage scenario achieved the highest peak performance depended on the complexity level *K* (Figure 9). At the additive case of *K* = 1, all strategies achieved very similar peak performances close to the theoretical maximum of 2/3. Until *K* = 8, the highest peak performances were reached with proportional usage of selected inbred lines. For *K* > 8, disproportional use of inbred lines resulted in the highest peak performances. Balanced use generally resulted in the lowest peak performance, except for *K* < 4, where this strategy was slightly ahead of the disproportional usage strategy. The differences between the strategies tended to increase with *K*.

**Figure 9.**
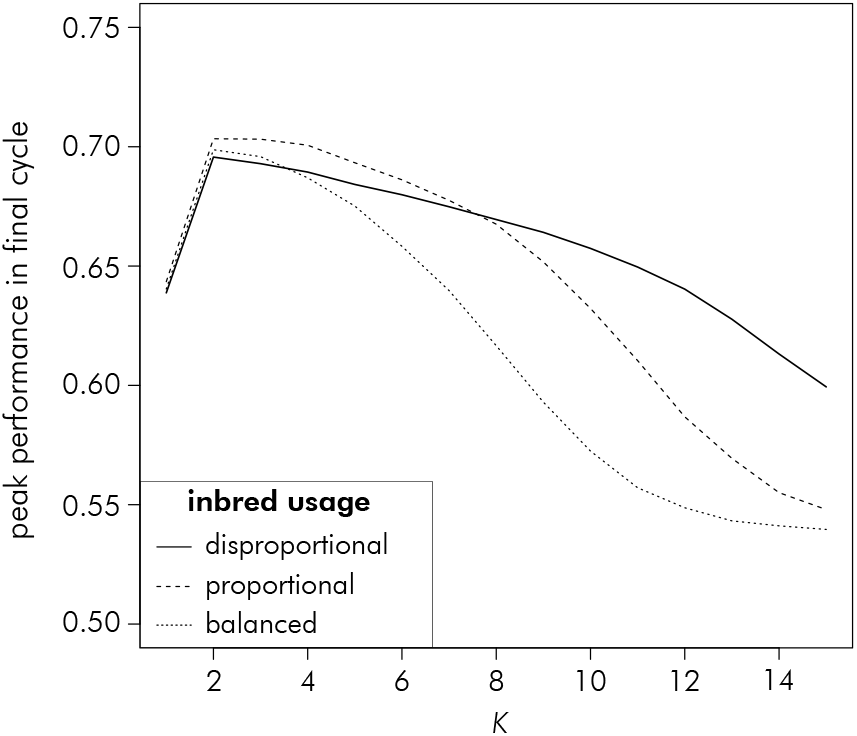
Relationship between the *NK* model complexity parameter *K* and peak genetic performance in the last cycle for the strategies explored in the the inbred usage theme.

The cycle over cycle increase in peak performance was initially higher the more disproportional the use of the inbreds (Figures 10A, B, C). However, except for the highest level of complexity, this did not result in the highest maximum performance for this scenario, because the increase started to level off in the last five to ten cycles. Scenarios with proportional and balanced use of inbreds therefore had the highest peak performance at *K* = 1, though the differences were small. At the intermediate level of *K* = 6, the disproportional use scenario was overtaken by the proportional use scenario in the last cycles. The differences between these two were small, however. Finally, at *K* = 15, only the disproportional use scenario achieved a sizable increase in peak performance.

**Figure 10.**
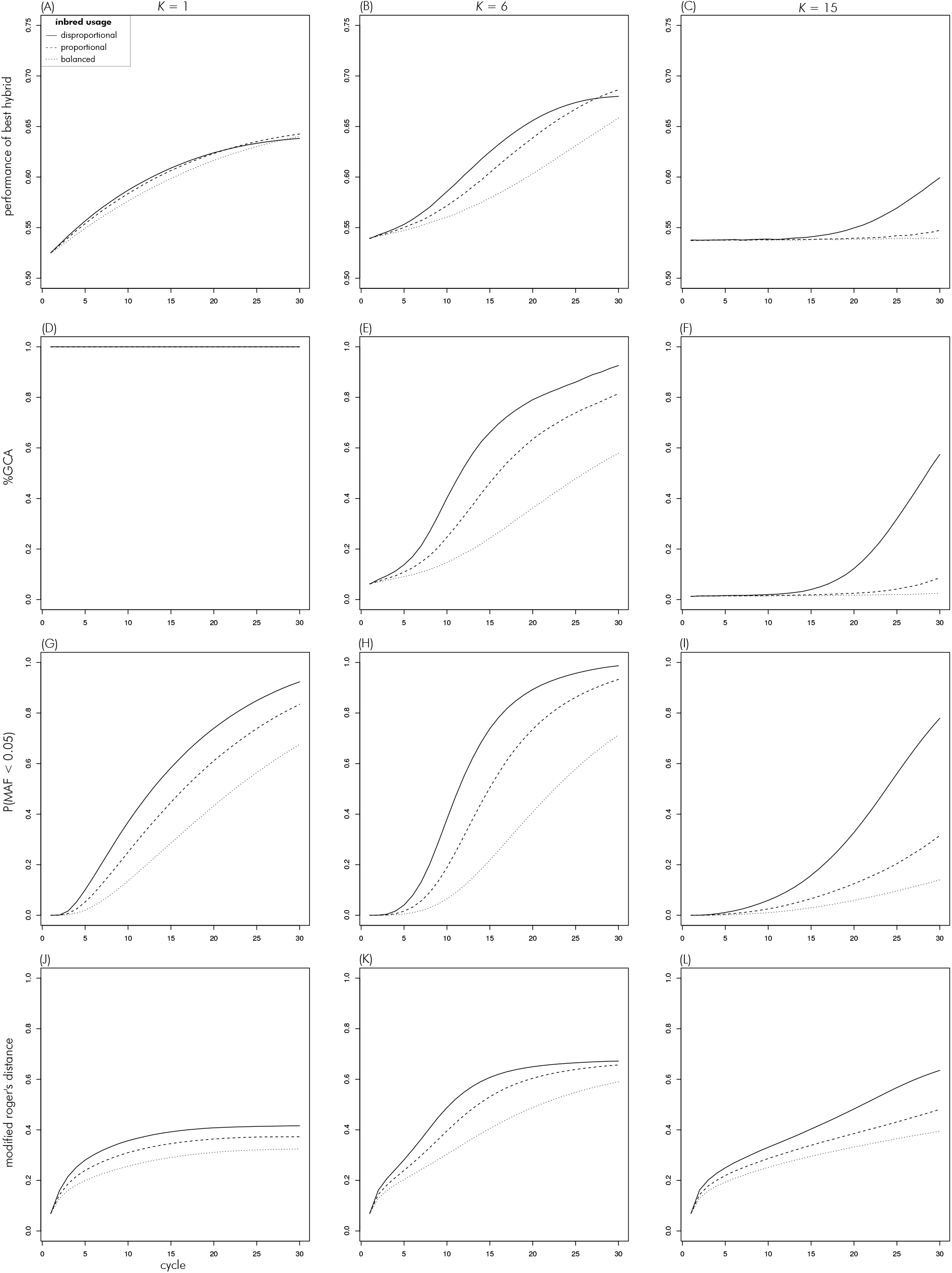
Evolution of metrics over cycles in the inbred usage theme for scenarios with *K* of 1 (left column), 6 (middle) and 15 (right).

At *K* = 1, %GCA stayed constant at one for all scenarios, as expected. At *K* = 6, %GCA increased most strongly for disproportional use, followed by proportional and balanced use (Figures 10D, E, F). Reaching above 90% for the former, and above 80% and 50% for the latter two, respectively. At *K* = 15, %GCA remained near zero for the balanced and proportional use scenarios throughout. For the disproportional use scenario, it remained at zero as well until cycle ten and increased from there to almost 60%.

For all strategies and values of *K*, the percent of loci with a MAF < 0.05 increased from its initial value of zero (Figures 10G, H, I). The increase over cycles was strongest for disproportional inbred use, for which it reached close to 100% at *K* of 1 and 6. The proportional usage strategy had the second strongest increase and the balanced strategy the weakest. As was the case in the decentrality theme, the differences between the strategies tended to increase with *K*. At the highest level of *K* = 15, the proportional and balanced strategies stayed below 40%, whereas the disproportional usage strategy reached close to 80%.

The modified Rogers distance between heterotic groups increased over cycles in all scenarios (Figures 10J, K, L). Throughout it was highest for disproportional use, followed by proportional and balanced use. Overall, the distance was greatest for the intermediate complexity level of *K* = 6.

*N*_*e*_, again reported only for cycle 15 at *K* = 7, was 23.2, 31.3 and 44.6, for the disproportional, proportional and balanced usage scenarios, respectively.

## Discussion

The objective of this study was to explore properties of the historically grown structure of commercial hybrid breeding programs, particularly in maize, and aid the understanding of why these structures successfully generated significant amounts of genetic gain in the past and thereby impact global food security. The infinitesimal framework (Barton et al., 2017), in which traits are described as the sum of a large number of genes all having additive, context independent effects of similar magnitude, is our starting point. However, we seek extensions to account for the empirical observations that a) there are results observed from operating a long-term breeding effort that are not consistent with or not easily explained within the infinitesimal model framework (Rasmusson and Phillips, 1997) and b) reflect the reality of a highly complex trait biology (Hammer et al., 2006). Therefore, we are motivated to consider the influence of complexity of trait genetic architecture on breeding strategies from the perspective of a long-term commercial breeding program (Duvick et al., 2004; Cooper et al., 2014).

### Emergence of additivity

As a representation of genetic complexity we chose the *NK* model framework developed by Kauffman (1993), which allows exploration of the full continuum from complete additivity to deep and almost intractable genetic complexity. For reference, the *NK* models used in this study, corresponded to the Mount Fuji landscape at *K* = 1 (Figure 1) and to the ‘Alps’ landscape from *K* = 2 to *K* of 7 or 8. After this the genetic models transitioned from the multi-peaked but correlated ‘Alps’ landscape to the uncorrelated landscape represented by the ‘Dunes’ metaphor (Figures 1 and 2).

In complex genetic landscapes, additive genetic variance, the sine qua non of genetic gain, is not a constant factor of trait biology (i.e., deducible from the molecular properties of genes) but rather emerging from the interplay of biology and natural or artificial properties of population structure (Wade, 2002; Cooper et al., 2005). In particular, additivity emerges in response to a constraining of the dimensionality of genetic space, or, in other words, by limiting genetic diversity. In practice, such constraints in dimensionality are achieved through fixation or near fixation of genes (Wade, 2002; Hill et al., 2008). This process is illustrated in Figure 11 for a simple epistatic network consisting of two genes. Thus, as genetic complexity increases, the breeder needs practical ways to reduce this complexity to a manageable level that allows genetic progress. This study explored two particular practical approaches that have been adopted within commercial hybrid breeding, particularly in maize. With the availability of genomics and novel thinking about genetic complexity, we can now study the genetic implications of these practical approaches, many of which were devised and adopted prior to the availability of a theoretical and empirical framework to study their effects.

**Figure 11.**
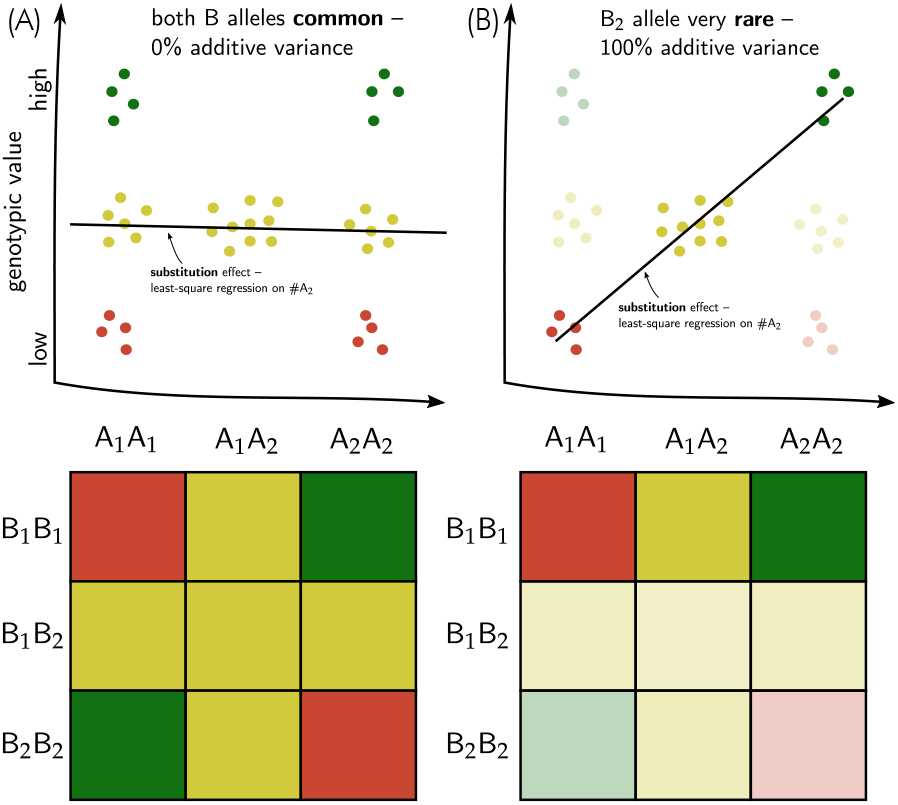
Illustration of a two-locus epistatic network in which neither the A nor the B locus exhibit any additive variation when all alleles are common (A) but collapses to a perfectly additive system in which the substitution effect of the A locus explains 100% of the variation when the B_2_ allele becomes rare (B). Note that the substitution effect of the A locus would be reversed in sign when the B_1_ allele became rare instead. Colours green, yellow and red represent high, intermediate and low phenotypic values, respectively.

Two processes in particular accelerate such constrainment, namely the creation of population bottlenecks and the subdivision of larger populations into more or less independent ‘demes’ (Katz and Young, 1975; Goodnight, 1995). Equivalent processes in the context of plant breeding programs are the degree of connectivity between breeding programs and the relative use of superior inbred lines in breeding crosses for producing the next generation, both ‘themes’ were explored in this study.

### Decentrality theme

Classical quantitative genetics infinitesimal theory was used to design and optimize commercial hybrid breeding strategies, in combination with em pirical experience of what worked and what did not (Hallauer et al., 2010). Yet, even though the infinitesimal model implies optimality of a single, homogenous population, there were discussions about the relative merits of large centralized vs. decentralized breeding programs early on (Baker and Curnow, 1969). Later on, Podlich and Cooper (1999) explored this problem on the basis of Sewall Wright’s *shifting balance theory*, (Wright, 1931, 1977; Wade and Goodnight, 1998). The shifting balance theory describes an evolutionary process in which genetic drift resulting from population subdivision enables random movements across genetic space (i.e., against selection gradients) and also converts epistatic to additive genetic variation through constraining genetic space as described above. This then enables local adaptation in complex genetic landscapes which is followed by differential migration from higher to lower performing sub-populations and thus ‘spreading’ of superior gene complexes across the whole *meta population*.

While this theory remains controversial as a model of natural evolution (Coyne et al., 1997), there are remarkable similarities between meta populations in the context of the shifting balance theory and the population structure of large commercial breeding operations. The latter also do not operate as a centralized unit but rather as a decentralized network of smaller programs with the most successful germplasm being shared across (Cooper et al., 2014). The same seems to be the case at the industry level, with the major commercial breeding operations being based on unique and distinct germplasm backgrounds, with only occasional exchange of elite material, e.g., through ex-PVP lines (Mikel and Dudley, 2006; White et al., 2020). As a result of this decentralization, plant breeding programs are also characterized by having low effective population sizes (Cowling, 2013), which makes them more susceptible to genetic drift.

Here, we expanded on the work of Baker and Curnow (1969) and Podlich and Cooper (1999) by exploring breeding population structures with differing degrees of decentrality (Figure 4), ranging from a large centralized program with high *N*_*e*_ to a isolated set of smaller programs with low *N*_*e*_, with a series of scenarios with decentralized but connected programs with *N*_*e*_ values in between these two extremes. We indeed found that strategies resulting in low within program *N*_*e*_ through increased decentralization and isolation became increasingly superior in terms of peak hybrid performance, as genetic complexity *K* increased, while a centralized strategy with high *N*_*e*_ was superior in less complex landscapes. These results thus confirm the findings of Podlich and Cooper (1999) that a decentralized search strategy is superior in complex genetic landscapes. Increasing isolation and decentralization and the associated *N*_*e*_ reduction led to quicker increases over cycles and higher overall values of of %GCA (Figure 8). At the highest levels of complexity, only complete isolation generated amounts of GCA variation sufficient for making genetic improvements cycle over cycle. The better ability to expose additivity in the form of GCA variation of the more isolated and decentralized strategies was expected as per the discussion at the beginning of this section outlining the relationship between amounts of additive variation and constrainment of genetic space. This explains the clear advantages in terms of genetic peak performance of the isolated strategy at the highest levels of *K*.

The corollary of the constrainment of genetic space of course is a more rapid decline in genetic diversity and susceptibility to genetic drift, which ultimately limits the selection potential of the programs. Indeed, for values of *K* below eight, which marked the switch from an uncorrelated to multi-peaked but correlated genetic landscape (Figure 2), decentralized programs with increasing rates of germplasm exchange became superior. Accordingly, having a large centralized program became the optimal strategy at lower values of *K*. Here, the genetic landscape was simple enough to not require severe constrainment of genetic space to expose sufficient amounts of GCA variation. The genetic drift experienced by small, isolated programs then unnecessarily led to the fixation of unfavourable alleles. This was most apparent at *K* = 1 where all variation is additive by definition and a decentralized strategy is not expected to have any advantage (Rathie and Nicholas, 1980). Here the isolated strategy led to fixation of almost all loci from cycle twenty onward and to a stalling of genetic gain significantly below the theoretically achievable maximum (Figure 8).

The establishment of genetically divergent heterotic groups has always been a central tenant of hybrid breeding (Melchinger and Gumber, 1998). Originally, optimal exploitation of heterosis was the main driver of their establishment (East, 1936). Later, however, maximization of GCA vs. SCA variation was identified as an import secondary feature of heterotic groups (Melchinger and Gumber, 1998). While this is well established for dominant gene action (Reif et al., 2007; Fischer et al., 2009), there are also indications for the conservation of favourable epistatic patterns that are disrupted when lines from different heterotic groups are recombined (Bernardo, 2001). Often, heterotic groups are established from populations that evolved in isolation for a long time. One of the best examples for this is the Dent by Flint heterotic pattern in maize which is prevalent in Central Europe and is comprised of populations that evolved in separation for centuries (Rebourg et al., 2003). Heterotic groups are thus a different and additional form of constrainment of genetic space through historically grown genetic isolation. In our simulations, the different heterotic groups were originated from the same base population, yet we still observed a significant degree of genetic differentiation evolve over cycles (Figure 8), as expected in recurrent, reciprocal selection regimes (Labate et al., 1999; Longin et al., 2013). A portion of this differentiation can be attributed to genetic drift (Gerke et al., 2015), as evidenced by the non-zero genetic distance at *K* = 1, where all effects are additive and increasing genetic differentiation between heterotic groups would have no effect on the proportion of GCA variance. However, the genetic differentiation was considerably higher for *K* > 1 (Figure 7), indicating that there indeed was a selection advantage to increased heterotic group divergence in complex genetic landscapes. This was particularly clear for the isolated scenario, where heterotic patterns could form uninterrupted within programs. For the centralized and decentralized strategies, the differentiation was maximal at lower values of *K*, because %GCA, and hence the effectiveness of recurrent, reciprocal selection, declined afterwards.

### Inbred usage theme

The history of North American and European maize germplasm can be described as a succession of key inbreds that were heavily used in breeding crosses and had a distinct and lasting impact on germplasm (Mikel and Dudley, 2006; Technow et al., 2014; White et al., 2020). Those inbreds owe their success either to the outstanding general combining ability relative to their peers at the time, such as was case for the important North American line B73 (Mikel and Dudley, 2006) or their unique adaptation to specific climatic conditions, such as the European Flint lines *F2* and *F7* (Messmer et al., 1992; Böhm et al., 2014). The highly disproportionate importance of successful inbreds led to a significant reduction in genetic diversity (Rasmusson and Phillips, 1997; White et al., 2020), particularly relative to the source populations from which they were derived (Böhm et al., 2017). However, this constrainment also might be responsible for the emergence of additive genetic variation from complex gene action through the so called founder or bottleneck effect (Goodnight, 1988; Cheverud and Routman, 1996; Naciri-Graven and Goudet, 2003; van Heerwaarden et al., 2008). We indeed observed that %GCA increased faster over cycles and reached higher values overall the more uneven the use of selected parents in breeding crosses (Figure 10), with the exception of *K* = 1, where all variance is additive by definition. At the highest degrees of landscape complexity only disproportionate use of inbreds, resulting in very low *N*_*e*_, succeeded in generating amounts of %GCA sufficient for genetic improvements. Like in the decentrality theme, the higher values of %GCA of the disproportional use strategy translated into superior peak performances only at the values *K* > 8, i.e., after the landscape transitioned from multi-peaked but correlated to uncorrelated. Before that, balanced and particularly proportional use, both having higher *N*_*e*_, achieved superior peak performances.

### Maintenance of diversity

In our simulations, the constrainment of genetic space and reduction of *N*_*e*_ through decentralization and isolation or disproportionate use of inbred lines, while necessary for exposing additive genetic variation, led to a rapid fixation of alleles and a slowing of genetic gain in later cycles. This was partly a consequence of genetic drift caused by the low *N*_*e*_ (Cowling, 2013). However, the reduction of *N*_*e*_ was also caused in part by the effects of selection, particularly once the majority of the genetic variation was additive. This has not generally happened in commercial breeding programs, where genetic gain continues apace (Rasmusson and Phillips, 1997; Fischer et al., 2008; Duvick et al., 2004; Pfeiffer et al., 2019). Several factors that maintain diversity in practical programs were not included in the simulation model. For example, the simulation implicitly assumed that the environment and management conditions remained constant across all cycles, whereas both change more or less rapidly in reality. Changing selection environments imply changing selection targets and trajectories (Messina et al., 2011; Hammer et al., 2009), which reduce the pressure on particular alleles or allele complexes and thus slow or prevent fixation. Long-term selection experiments have shown that selection response can be maintained even in isolated and narrow populations (Dudley and Lambert, 2010; Durand et al., 2010, 2015). Several hypotheses were proposed for these surprising results, including epistasis (Carlborg et al., 2006), de novo genetic mutations, particularly when magnified through effects on epistatic complexes (Rasmusson and Phillips, 1997; Durand et al., 2010), creation of heritable epigenetic variation (Hauben et al., 2009), activity of transposable elements (Dubin et al., 2018), as well as the presence of ‘cryptic genetic variation’ through phenomena such as canalization (Gibson and Dworkin, 2004). Of these, only epistasis was present in our simulations. While highly speculative, these biological phenomena might explain the presence of abundant genetic variation and continued genetic gain in largely isolated and genetically narrow commercial plant breeding programs.

### Applications of the *NK* for plant breeding

The *NK* model, originally developed by the theoretical biologist Stuart Kauffman to study evolution in complex genetic landscapes, has found applications for modelling complex systems in disparate fields such as business administration (Csaszar, 2018), organizational learning theory (Lazer and Friedman, 2007), infrastructure design (Grove and Baumann, 2012) and physics (Qu et al., 2002). Following the example of (Podlich and Cooper, 1999), we here used the *NK* model to represent genetic complexity navigated by commercial hybrid breeding operations to study the effect of the degree of isolation between programs as well as the degree of imbalance in inbred usage, both key aspects of breeding strategies. We propose that this model can serve as an ideal starting point to study other aspects of hybrid breeding strategies. For example, Cooper and Podlich (2002) proposed an extension to the *NK* model that adds an environmental dimension and thus allows modelling concepts related to genotype by environment interaction (Cooper and DeLacy, 1994), yield stability (Piepho, 1998; Tollenaar and Lee, 2002), product placement (Messina et al., 2018) and the target population of environments (Comstock, 1977). These so called *E(NK)* models represent different environments or management practices through a series of more or less similar genetic landscapes. This of course adds considerable complexity to the already complex static landscapes studied here and poses interesting dilemmas. For example, rapidly exposing additive variation, e.g., through isolation, might be even more important than in static landscapes because local optima have to be exploited quickly before they disappear once the environment shifts, for example through changes in management such as the historical increases in plant population for commercial maize production (Hammer et al., 2009). However, retaining allelic diversity, which hampers the exposing of additivity, is required to enable renewed adaptation to the changed environmental conditions.

A high degree of genetic complexity also implies a high degree of context dependency of genetic effects. Observe, for example, that the additive substitution effect of the ‘A’ locus in Figure 11 changes sign when the B_1_ allele instead of the B_2_ allele becomes rare. This has consequences on the persistence of accuracy of estimated QTL effects and genomic prediction models and can be addressed through iteratively updating training populations for genetic model parameterization (Podlich et al., 2004; Wolc et al., 2016; Forneris et al., 2017). The *NK* model framework can help address questions about the frequency with which this has to happen and whether data from previous generations can be used. Recently, approaches were proposed that attempt to capture those context dependencies through biological models representing the interdependencies underlying the traits of interest (Technow et al., 2015). Such models are only approximations of the true biological complexity. However, Cooper et al. (2005), using the *NK* framework, have shown that even incomplete knowledge of biological networks can improve predictability of genetic effects and genetic gain. The context dependency of genetic effects, i.e., the effects being neither universally positive or negative (Wade, 2002), also has implications on innovative proposals for using CRISPR-Cas9 gene editing (Jaganathan et al., 2018; Gao et al., 2020) to either target recombination to create superior hypothetical linkage groups (Brandariz and Bernardo, 2019) or even the large scale “editing away” of deleterious mutations (Wallace et al., 2018). Finally, this framework might help devise strategies for the efficient introduction of exotic or ancient germplasm (Yu et al., 2016; Böhm et al., 2017), which evolved not just in a very different environmental, but also a different genetic context from the current elite breeding germplasm.

### Back to the future

The structure of commercial plant breeding programs, particularly in major crops like maize, is characterized by a large degree of decentralization with exchange of successful germplasm within companies (Cooper et al., 2014), while isolation is the norm among companies (Mikel and Dudley, 2006). Plant breeders further have a tendency for relying on only a small set of elite inbred lines for producing the next generation (Rasmusson and Phillips, 1997), leading to a series of significant bottleneck events (White et al., 2020). All of these features lead to a drastically reduced effective population size and are not predicted to be promising strategies under the additive, infinitesimal model of quantitative genetics. Yet commercial hybrid breeding has delivered incredible amounts of genetic gain over the last century, and has thus contributed to food security and resource conservation (Duvick, 1999). Here we postulated that the described structure of plant breeding programs, with its constrainment of genetic space, is in fact necessary for enabling the exploration and exploitation of genetic variation in complex genetic landscapes and that the success of a breeding program is not only determined by its germplasm per se, but by the structures that allow it to evolve. The breeding program structures described here grew historically and we do not claim that it was set up with this intention. However, by doing “what worked”, breeders in preceding generations might have nonetheless been able to take advantage of the process described and postulated in this study. Understanding why these historic structures “worked” will be critical for designing breeding programs that can tackle the challenges of the century ahead.

