## Supplemental File S1 for "Back to the future: Implications of genetic complexity for hybrid breeding strategies"

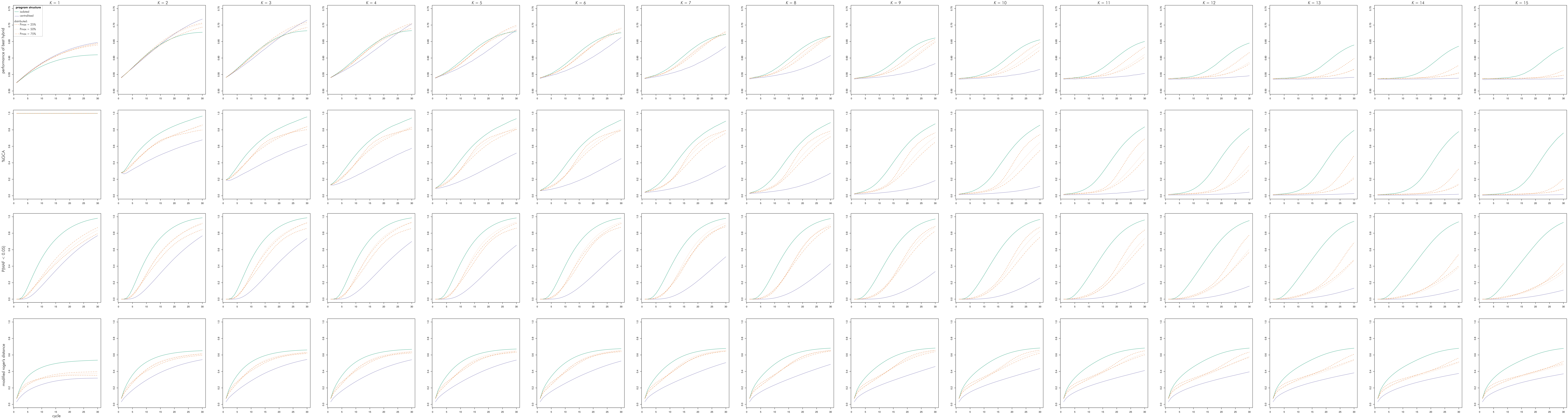


**Figure S1** Evolution of metrics over cycles in the decentrality theme for all values of complexity parameter *K* (columns)
