## Supplemental File S2 for "Back to the future: Implications of genetic complexity for hybrid breeding strategies"

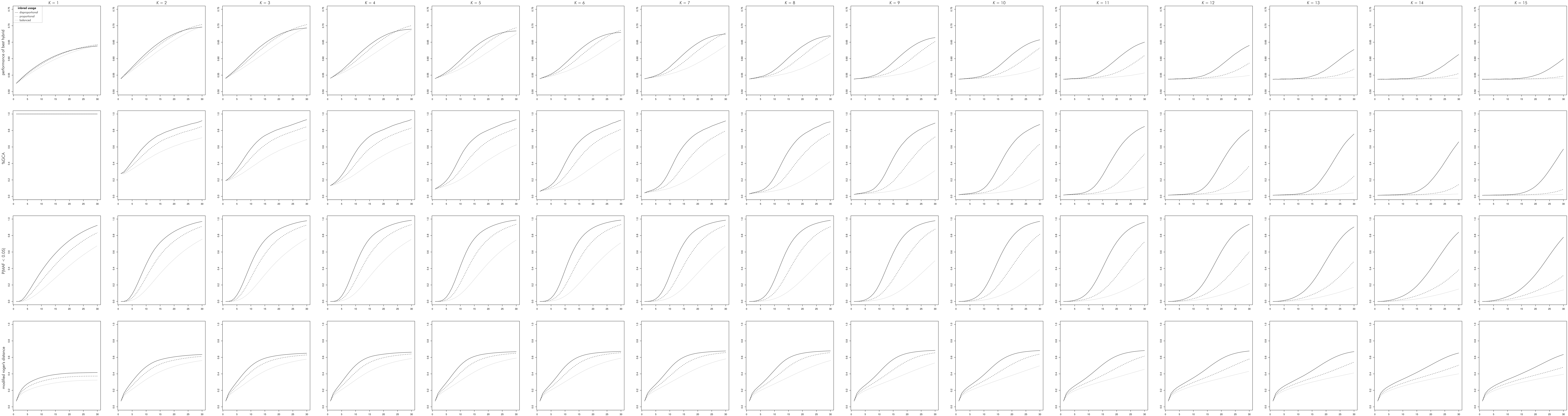


**Figure S2** Evolution of metrics over cycles in the inbred usage theme for all values of complexity parameter $K$ (columns)
